# APC/C-Vihar regulates centrosome activity and stability in the *Drosophila* germline

**DOI:** 10.1101/202465

**Authors:** A. L. Braun, F. Meghini, G. Villa-Fombuena, M. Guermont, E. Fernandez-Martinez, D.M. Glover, M. D. Martín-Bermudo, A. González-Reyes, Y. Kimata

## Abstract

A universal feature of metazoan reproduction is the elimination of the maternal centrosomes prior to the end of oogenesis. In animals that have a syncytial cyst stage of oocyte development, including *Drosophila* and mouse, the germline centrosomes undergo a migration to all reside within the oocyte. However, the functional significance of centrosome transport within the female germline and the mechanism orchestrating this event are still a mystery. The Anaphase Promoting Complex/Cyclosome (APC/C) is a multi-subunit ubiquitin ligase (E3) that temporally regulates progression of the cell cycle as well as the centrosome cycle. By altering the negative regulation of the cooperating ubiquitin conjugating enzyme (E2), Vihar/Ube2c, we show that temporal control of APC/C activity ensures centrosome stability and migration during early *Drosophila* oogenesis. When there is perduring APC/C activity, Polo kinase is precociously targeted for destruction, which results in centriole instability and decreased centrosome transport to the oocyte. We show that decreased centrosome transport correlates with a decreased accumulation of pericentriolar material (PCM) proteins on the oocyte nucleus, which results in a weakening of the structural integrity of the egg chamber and loss of oocyte fate – the overall consequence being a reduction in female fertility. Considering the conserved roles of the APC/C and Polo kinase throughout the animal kingdom and the fact that many animals have a syncytial stage of egg development, our results provide insight into the general necessity of gametic centrosome transport for female fertility.

## Introduction

Centrosomes function as microtubule organizing centers (MTOC) in many animal cells, they consist of a pair of centrioles and an amorphous proteinaceous matrix, the PCM, which is responsible for the spatial concentration of microtubule nucleating proteins. As cells divide, the centrosomes focus the mitotic spindle and can influence cell fate decisions (Lattao et al., 2017). In animals that need sperm to activate the egg, the centrioles are thought to be eliminated from the egg in order to prevent parthenogenesis (Manandhar et al., 2005). However, in *Drosophila melanogaster*, where the egg is activated prior to fertilization, forced centriole retention results in meiotic defects and subsequent sterility (Pimenta-Marques et al., 2016). Conversely, premature centriole elimination is thought to have no effect (Stevens et al., 2007). *Drosophila* egg development proceeds in 14 stages (S) that are delineated by morphological changes (Fig. 1a) (Roth and Lynch, 2009). From a 16-cell syncytial cyst, one cell – eventually marked by persistent synaptonemal complexes, centrosome accumulation, and fate determinant localization (Huynh and St Johnston, 2004) – ultimately becomes the oocyte, whilst the other cells take on a nurse cell fate. The oocyte arrests in meiosis I, whereas nurse cells undergo several rounds of endoreplication and function to provide the oocyte with all of the materials necessary to carry out embryogenesis. The purpose of the centrosomes throughout oogenesis is unknown; however, during later stages they may retain their microtubule organizing functionality to help maintain the cytoskeleton of the egg chamber (Grieder et al., 2000; Januschke et al., 2006; Theurkauf et al., 1993; Tissot et al.,. Since the correct microtubule organization is essential for the proper establishment of the primary and secondary axes of the *Drosophila* body plan (Roth and Lynch, 2009), it is important to establish if the centrosomes aid in microtubule organization during oogenesis.

**Figure 1:**
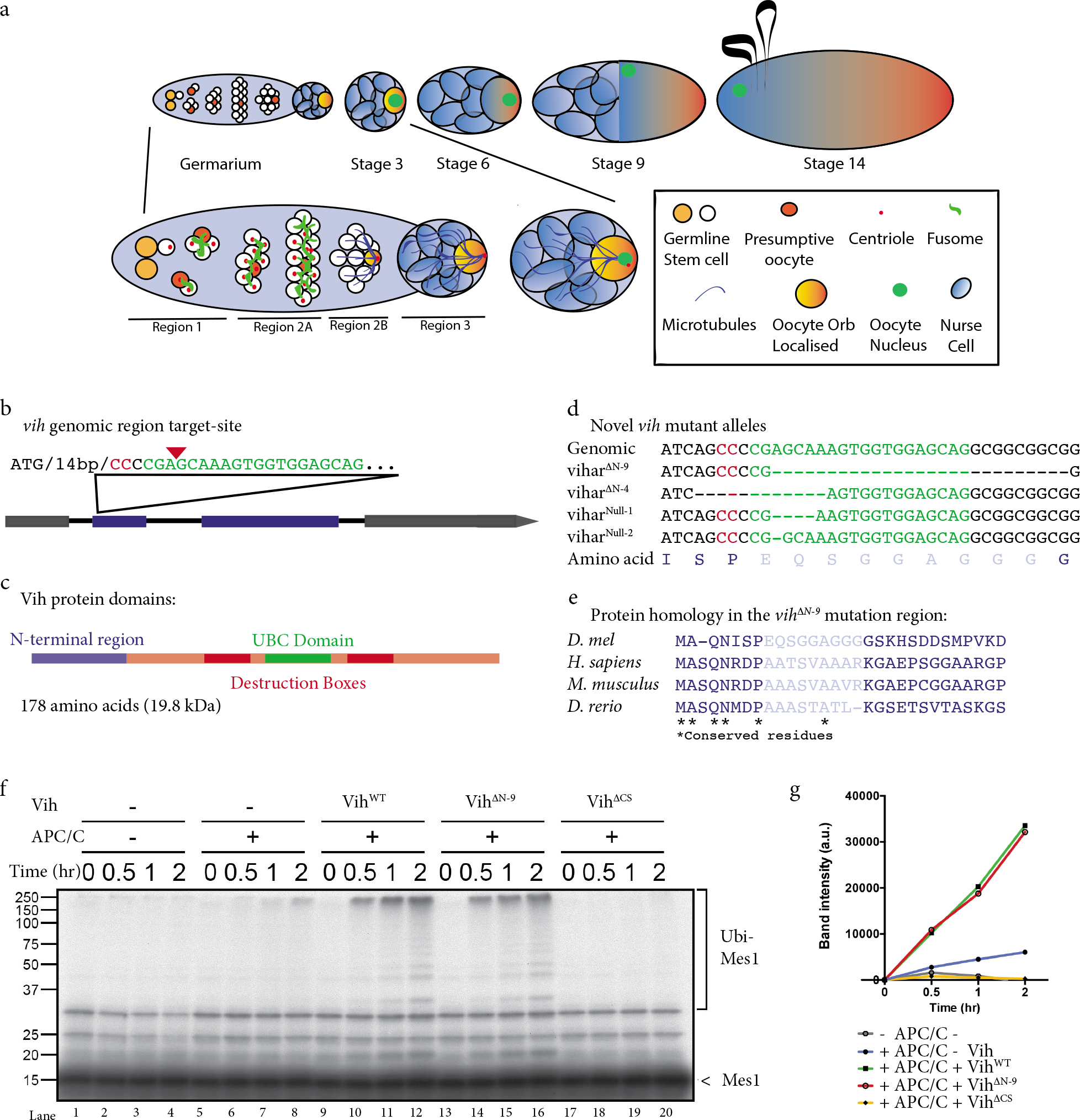
Egg chamber development schematic, vih allele design and Vih^ΔN-9^ working with the APC/C in vitro **a)** Schematic egg chamber development. **b)** CRISPR/Cas9 genomically engineered gRNA target-site design within the coding region of the gene. The red arrow denotes the cut site, the red residues denote the protospacer adjacent motif (PAM) sequence, and the green residues denote the gRNA complementary sequence. **c)** Protein schematic and sequence for Vih. **d)** Four examples of vih alleles that were created. **e)** Protein homology in vertebrates for the region that we altered in our *vih^ΔN^* alleles. **f)** Drosophila reconstituted in vitro ubiquitination assay using Mes1 as a substrate: lanes 1-9 with no Vih, lanes 9-12 with Vih^WT^, lanes 13-16 with Vih^ΔN-9^, and lanes 17-20 with Vih^ΔCS^. **g)** Ubiquitination assay quantification.

### CRISPR/Cas9 generated vihar/Ube2c alleles alter APC/C activity

In most cells, the centrosome microtubule organizing activity and duplication process are tightly coupled to cell cycle progression (Prosser et al., 2012). This coupling is, in part, mediated by the APC/C. The APC/C targets many cell cycle proteins (including mitotic cyclins) as well as proteins that regulate centriole biogenesis (such as human Sas6 and STIL) for destruction by the proteasome within specific time windows during the cell cycle (Davey and Morgan, 2016). Therefore, to investigate the role of centrosome activity during oogenesis, we decided to perturb APC/C activity by targeting the highly-conserved APC/C-specific E2, Vihar (Vih; Ube2c in human) (Mathe et al., 2004; Yu et al., 1996). In the fly, Vih is necessary for Cyclin B (CycB) degradation by the APC/C and its loss-of-function results in mitotic arrest (Mathe et al., 2004). In both humans and flies, Vih/Ube2c is localized to the centrosome and its cellular levels oscillate during the cell cycle, peaking during mitosis and decreasing into interphase (Mathe et al., 2004; Rape and Kirschner, 2004; van Ree et al., 2010), suggesting a critical role of Vih/Ube2C in spatiotemporal regulation of APC/C activity. Using CRISPR/Cas9 genomic engineering, we specifically targeted the 27-amino acid N-terminus of the Vih protein creating a series of *vih* alleles (Fig. 1b, c; see Methods for detailed characterization). The N-terminal domain has been shown to decrease auto-inhibition of Ube2c in human cell lines (Summers et al., 2008), resulting in sustained APC/C-Ube2c activity. Two homozygous lethal *vih* null alleles (*vih^Null^*) were created by frame-shifts (Fig. 1d), which had similar cell cycle phenotypes in the larval neuroblasts to previously reported null alleles; i.e., chromosomes were disorganized and there were supernumerary centrosomes (Extended Data Fig. 1a) (Mathe et al., 2004). In-frame deletions (*vih^ΔN^*) were also created, covering residues that are highly conserved in vertebrates (Fig. 1d, e). The reduced protein sizes of the two largest deletions (27 bp for *vih^ΔN-9^* and 12 bp for *vih^ΔN-4^*, Fig. 1d), were confirmed (Extended Data Fig. 1b).

We developed a reconstituted in vitro ubiquitination assay using the *Drosophila* APC/C to assess APC/C-Vih^ΔN-9^ (larger deletion) ubiquitination activity on the model APC/C substrate Mes1 (see Methods). As expected, Mes1 was ubiquitinated by the APC/C in the presence of wildtype Vih^WT^, but not in the presence of a catalytically inactive version, Vih^ΔCS^ (Fig. 1f, g). The APC/C achieved a similar to wildtype level of ubiquitination in the presence of Vih^ΔN-9^(Fig. 1f, g), indicating Vih^ΔN-9^ remains catalytically active. This finding is in agreement with previously published results where human Ube2c lacking the entire N-terminal domain performed equally as well as the full-length protein in in vitro ubiquitination and destruction assays (Summers et al., 2008).

### Egg chamber development is disrupted in vih^ΔN^ mutant females

In the germarium, the cystoblast undergoes four rounds of mitotic division with incomplete cytokinesis to create a 16-cell syncytial cyst, where cells are connected by intracellular bridges called ring canals (Huynh and St Johnston, 2004). Since the APC/C-Vih complex is necessary for mitotic progression, we first confirmed that there were 16 cells in the *vih^ΔN-4^* and *vih^ΔN-9^* germline cysts, indicating that the N-terminal amino acid deletions did not block cell division (Fig. 2a). The cysts appeared normal until S3-4, where we noticed cortical F-actin in germline cells was collapsing between two or more of the nurse cells, instead of localizing to cell peripheries, and as the number of membrane that collapsed increased the membranes eventually formed abnormal clumps with actin (in 45% of *vih^ΔN-9-/+^* egg chambers and 16% of *vih^ΔN-9-/+^*; Fig. 2a). The same phenotype was found at lower frequencies in *vih^ΔN-4^* egg chambers and never observed in control egg chambers (Fig. 2a), therefore we focused our work on the allele with the larger deletion. Since this “membrane phenotype” (as it will be termed hereafter) resembles that of mutations resulting in a loss of membrane integrity (Coutelis and Ephrussi, 2007; Tan et al., 2014), with time-lapse imaging we confirmed that the nurse cell membranes were rupturing instead of slowly disappearing, indicating that a physical force causes the membranes to break (Extended Data Fig. 1c, Movies 1-2).

**Figure 2:**
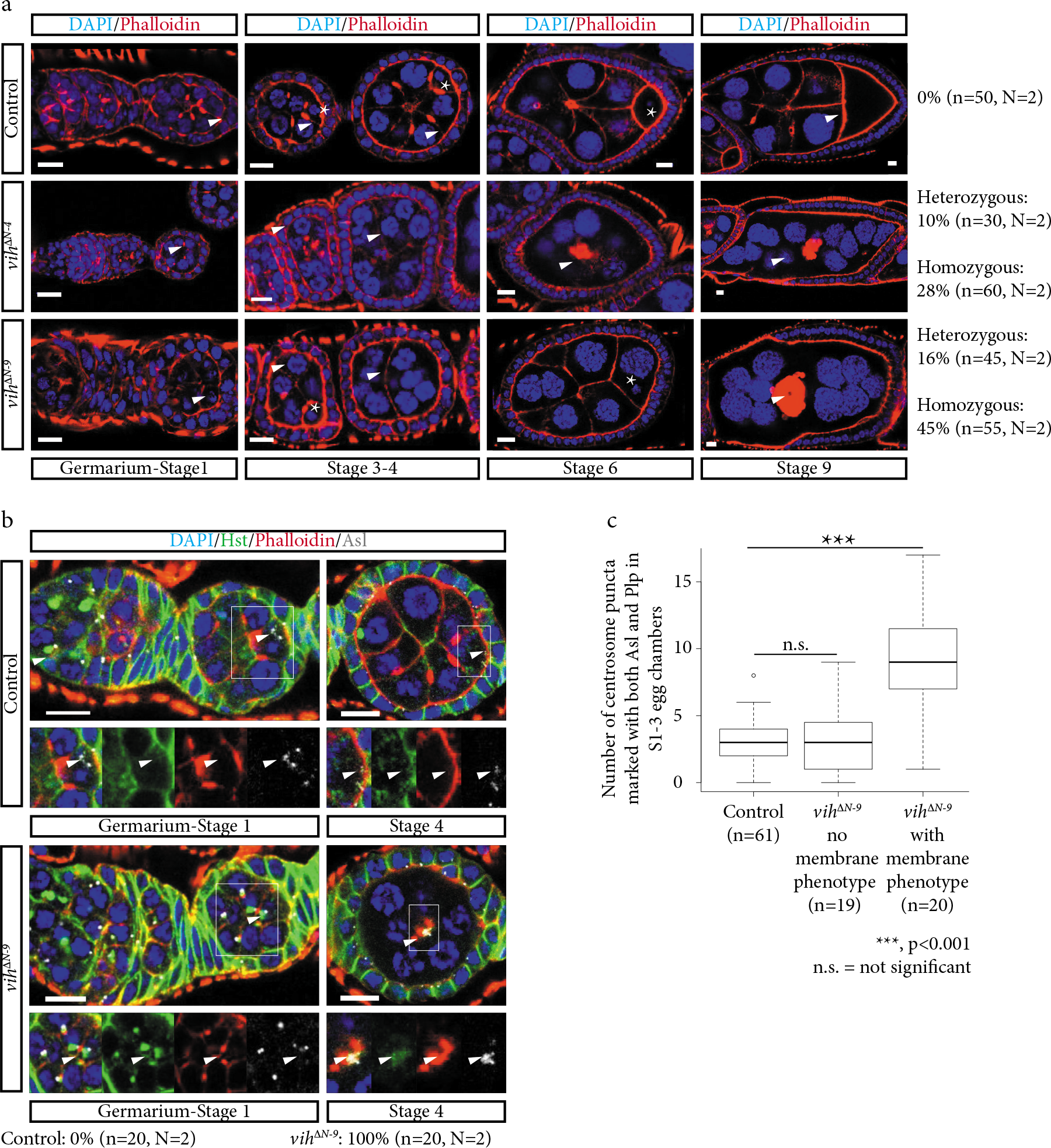
*vih^ΔN-9^* phenotype and centrosomes in the egg chamber **a)** Development time-series of phenotypic presentation for control, *vih^ΔN-4^*, and *vih^ΔN-9^* egg chambers, stained for DNA (DAPI) in blue and F-actin (Phalloidin) in red (arrows indicate the membranes and asterisks the oocytes). **b)** Germarium-S1 and S4 control and *vih^ΔN-9^* egg chambers stained for DNA (DAPI) in blue, fusome (Hts) in green, and F-actin (Phalloidin) in red, and centrioles (Asl) in white (arrows indicate centrioles). c) Quantification of ectopic centrosome puncta co-stained with Asl and Plp in S1-3 control and *vih^ΔN-9^* egg chambers, significance determined using a Welch’s t-test for the control compared with the phenotype (p=5.0×10^−6^) and with Wikcoxon signed-rank test with the control to no phenotype (p>2×10^−1^). All magnification bars equal to 10 μm.

### Centrosome transport and stability is compromised in vih^ΔN-9^ egg chambers

During the mitotic divisions that give rise to germline cysts, the centrosomes are associated with a temporary structure called the fusome, which forms a branching network that passes through each ring canal in the cyst (Fig. 1a) (Huynh and St Johnston, 2004). After mitotic exit, the centrosomes undergo intercellular transport from the nurse cells to the oocyte^2^. This process is dependent on Dynein, stable microtubules, and the fusome, but is independent of fate determinant localization (Bolívar et al., 2001; Huynh and St Johnston, 2004; Mahowald and Strassheim, 1970; Röper and Brown, 2004). In control egg chambers the majority of centrosomes are found in one cell, the future oocyte, where they subsequently localize to the posterior (Fig. 2b, c). While in S1-3 *vih^ΔN-9^* ovarioles that did not contain egg chambers displaying the membrane phenotype, an average of 3 centrosome puncta were found within the nurse cells, similar to controls (Fig. 2c). However, in egg chambers of *vih^ΔN-9^* ovarioles that exhibited the membrane phenotype at later stages, an average of 9 centrosome puncta are present within the nurse cells, often located near the ring canals (Fig. 2b, c). At later stages (S4-8), centrosomes were associated with the clumps of F-actin in the middle of the egg chambers (Fig. 2b). This failure in centrosome migration is not due to a disruption of the fusome, as fusome morphology in *vih^ΔN-9^* germaria appeared normal (Fig. 2b and Extended Data 2a). The tight correlation between the membrane and early centrosome transportation phenotypes suggests that both are linked.

During late oogenesis a cluster of oocyte nucleus-associated centrosomes is present until S12 in control egg chambers (Extended Data Fig. 2b) (Pimenta-Marques et al., 2016). However, in *vih^ΔN-9^* egg chambers the number of germline centrosomes were reduced after S6 in all *vih^ΔN-9^* egg chambers with the membrane phenotype and were virtually undetectable after S9 (Extended Data Fig. 2b), suggesting compromised centrosome integrity in *vih^ΔN-9^* egg chambers. To determine if decreased centrosomes stability contributes to the centrosome transport/membrane phenotypes, we tested whether further destabilizing the centrioles could increase the occurrence of centrosome transport failure and membrane phenotypes in *vih^ΔN-9^*egg chambers. We recombined *vih^ΔN-9^* with a *Sas4* mutant allele (*Sas4^s2214^*) expecting to further destabilize the centrosomes, since Sas4 is a core structural centriole component that is necessary for centriole duplication and may be required mitotic PCM recruitment (Lattao et al., 2017). We observed no centrosome transport defects in *Sas4^s2214^* heterozygous egg chambers alone (Extended Data Fig. 3a). However, we found that halving the dosage of Sas4 in homozygous *vih^ΔN-9^* egg chambers significantly increased the penetrance of the centrosome transport and membrane phenotypes from 45% (Fig. 2a) to 73% (Extended Data Fig. 3a, b). This result suggests that centriole instability is, at least in part, contributing to the centrosome transport failure and the subsequent membrane phenotype.

**Figure 3:**
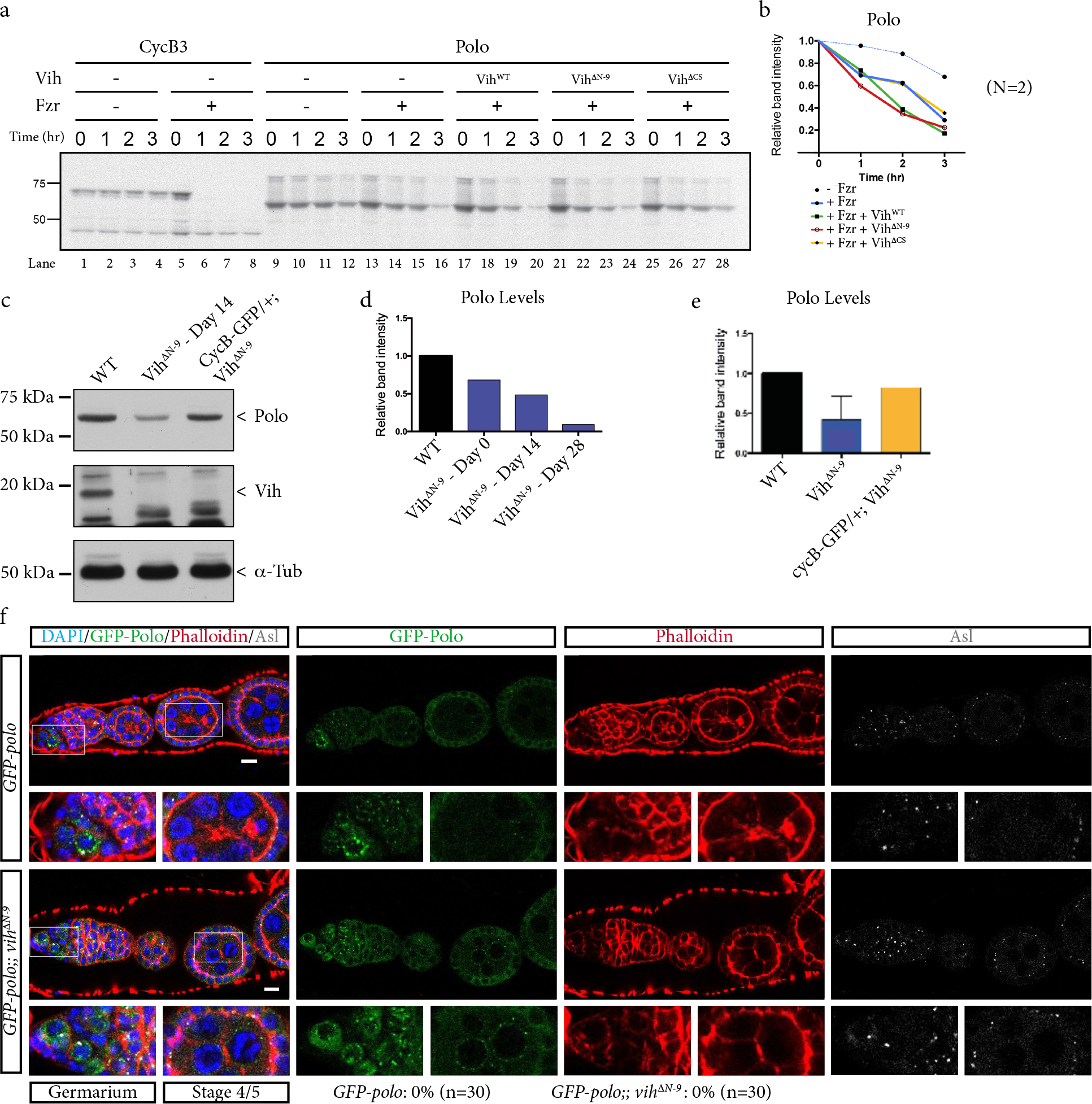
Polo kinase is a target of the APC/C in vitro and in vivo **a)** In vitro *Drosophila* destruction assay of Polo. Lanes 1-8 control CycB3 destruction. Lanes 9-28 Polo destruction: lanes 9-16 with no Vih, lanes 17-20 with Vih^WT^, lanes 21-24 with Vih^ΔN-9^, and lanes 24-28 with Vih^ΔCS^. **b)** Polo destruction quantification. **c)** Western blot of control, *vih^ΔN-9^*, and *CycB-GFP/+; vih^ΔN-9^* ovary extracts 14 days after eclosion, stained with Vih and Polo antibodies. **d)** Quantification of western blot Polo protein band intensity for control and *vih^ΔN-9^* ovary extracts at 0, 14, 28 days after eclosion. **e)** Quantification of Polo protein band intensity from western blots for control, *vih^ΔN-9^* flies 0, 14, and 28 days after eclosion averaged, and *CycB-GFP/+; vih^ΔN-9^* ovary extracts. **f)** GFP-Polo in control and *vih^ΔN-9^* egg chambers stained for DNA (DAPI) in blue, GFP-Polo in green, F-actin (Phalloidin) in red, and centrioles (Asl) in white. All magnification bars equal to 10 μm.

### Polo kinase regulates centrosome stability and transport downstream of APC/C-Vih

Having discovered a link between APC/C activity and centrosome stability, we investigated which APC/C substrate could account for the centrosome transport failure. Our prime candidate was Polo kinase (Plk1 in human): it is required for PCM recruitment to the centrioles (Prosser et al., 2012), is a known target of the APC/C in other organisms (Lindon and Pines, 2004), and in mammals is involved in the uncoupling of the centrosome cycle from the cell cycle during ciliogenesis (Al Jord et al., 2017). Furthermore, it was recently shown that Polo activity is crucial for centrosome stability in the *Drosophila* egg chamber (Pimenta-Marques et al., 2016). Using an in vitro destruction assay where APC/C-dependent proteolysis can be recapitulated in *Xenopus laevis* egg extract (Yamano et al., 2009), we confirmed *Drosophila* Polo is an APC/C substrate (Fig. 3a, b). Similar to the control APC/C substrate Cyclin B3 (CycB3), Polo was degraded more rapidly in interphase extracts with purified Fizzy-related (Fzr), an APC/C co-activator (Fig. 3a, b); it is important to note that *Drosophila* Polo shows a slow degradation even without APC/C activity in *Xenopus laevis* egg extract, which may be caused by activity of other E3 enzymes in the egg extract (Giraldez et al., 2017). Moreover, the addition of recombinant Vih^WT^ or Vih^ΔN-9^, but not Vih^ΔCS^, further accelerated Polo degradation in vitro. These results confirm that Vih cooperates with APC/C-Fzr to degrade *Drosophila* Polo and that Vih^ΔN-9^ is catalytically active.

To determine whether Polo levels are altered in *vih^ΔN-9^* egg chambers, we carried out western blots of *vih^ΔN-9^* ovary extracts. We found reduced levels of endogenous Polo protein in the *vih^ΔN-9^* ovaries compared to wildtype (Fig. 3c-e), suggesting an increase in Polo degradation in *vih^ΔN-9^* egg chambers. Next, we set to determine if the membrane and centrosome transport phenotype observed in *vih^ΔN-9^* egg chambers could be due to reduced Polo levels. We tested whether increasing the cellular amount of Polo, by introducing a functional GFP-Polo transgene, could rescue the *vih^ΔN-9^* phenotypes. Indeed, we found that two copies of GFP-Polo were able to completely rescue the membrane and centrosome transport phenotypes in the *vih^ΔN-9^* germline (Fig. 3f and 2a). These data support the idea that the membrane and centrosome transport phenotypes characteristic of *vih^N-9^* egg chambers originates from reduced Polo levels.

### Centrosome destabilization is caused by ectopic APC/C-Vih activity

Since we observed reduced Polo levels in the *vih^ΔN-9^* ovaries and the N-terminus is required for negative regulation of Vih, *vih^ΔN-9^* is likely to induce sustained APC/C activity. We first confirmed that the endogenous levels of the known APC/C substrate, CycB, are reduced in *vih^ΔN-9^* ovaries (Extended Data Fig. 3c). We subsequently reasoned that halving the dose of an APC/C catalytic subunit encoded by the *morula* (*mr*) gene should ameliorate the *vih^ΔN-9^*phenotypes. We found that only 3% of *mr^1^/+; vih^ΔN-9^* egg chambers showed the membrane phenotype, compared to the 45% typical of *vih^ΔN-9^* single mutants (Extended Data Fig. 3d and 2a). These data suggest perduring APC/C-Vih activity in *vih^ΔN^* egg chambers.

It has been suggested that APC/C targets its substrates in a sequential manner, in part, based on the efficiency of Ube2c to initiate the ubiquitin chain formation in each substrate, with high-affinity substrates being targeted before low-affinity ones (Williamson et al., 2011). Hence, APC/C-Vih^ΔN-9^ could be precociously targeting one of its low-affinity substrates. Considering that Polo levels are reduced in *vih^ΔN-9^* ovaries, we tested if Polo can be stabilized by the ubiquitous expression of a high-affinity substrate CycB in a *vih^ΔN-9^* background. We found that CycB-GFP overexpression rescued Polo levels (Fig. 3c, e) as well as the *vih^ΔN-9^* membrane phenotype (*vih^ΔN-9^* 45% Fig. 2a, CycB-GFP; *vih^ΔN-9^* 3% Extended Data Fig. 3e). Altogether, these results identify Polo as the downstream, lower-affinity target of APC/C-Vih that causes the centrosome and membrane phenotypes in *vih^ΔN-9^* ovaries. Overall these results highlight the critical importance of the precise control of centrosome activity by the APC/C and Polo to ensure egg chamber integrity during oogenesis in *Drosophila*.

### Oocyte associated PCM is reduced when there is decreased centrosome transportation

Since Polo is required for PCM recruitment, we sought to determine if the PCM is also reduced in *vih^ΔN-9^* egg chambers. We stained for Pericentrin-like protein (Plp), a PCM component that aids in interphase centrosome microtubule nucleation (Lerit et al., 2015). Similar to previous findings, in controls Plp labelled centrosomes until late oogenesis (Pimenta-Marques et al., 2016). Surprisingly, we also observed strong endogenous Plp staining on the oocyte nucleus/germinal vesicle, appearing before S1 and starting to taper off between S4 and S6 (Fig. 4a, b). Although a similar localization is previously reported with GFP-PACT overexpression (Martinez-Campos et al., 2004), the localization of endogenous Plp protein has not been reported. The Plp reduction that we observed coincides with the previously identified time points of Polo levels decreasing,^5^ and the localization of the MTOC and oocyte nucleus to the posterior of the egg chamber (Huynh and St Johnston, 2004; Pimenta-Marques et al., 2016). In S1/3 *vih^ΔN-9^* egg chambers where there were ∼9 centrosome puncta in the nurse cells but the membrane phenotype is not yet apparent, the Plp staining on the oocyte nucleus was significantly reduced (Fig. 4a, b). At later stages when the membrane phenotype appeared, the Plp staining was strongly reduced or absent. Since Plp is an MTOC marker and the oocyte nucleus is a known microtubule organizer, the reduction of Plp levels in the oocyte indicates that the MTOC activity of the oocyte nucleus is weakened even before the membrane phenotype arises, which supports earlier theories that the centrosomes may need to be proximal to the oocyte nucleus in order to transfer their MTOC capabilities (Tavosanis and Gonzalez, 2003; Theurkauf et al., 1993), similar to what is observed during human myogenesis (Tassin et al., 1985).

**Figure 4:**
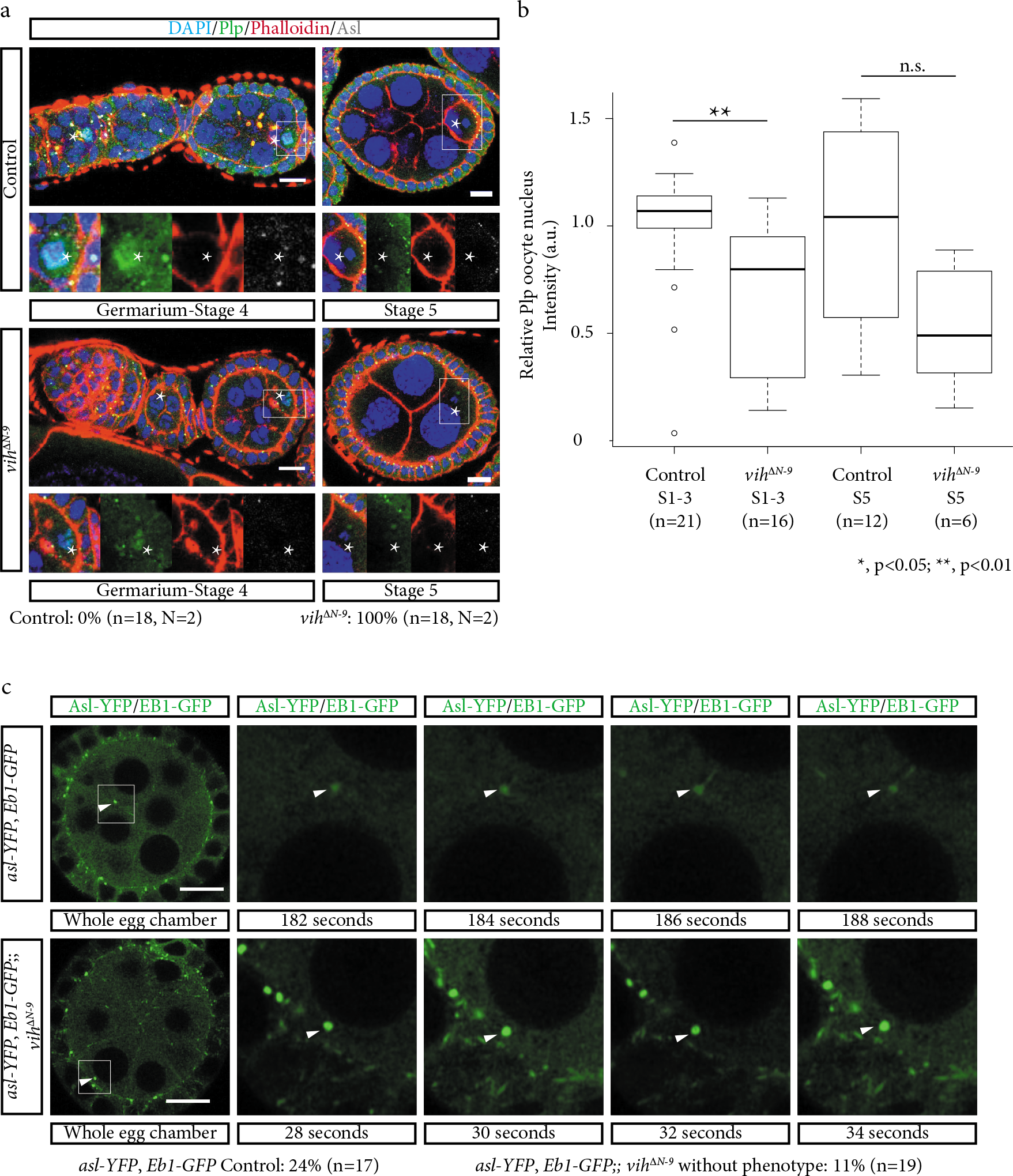
*vih^ΔN-9^* causes reduced PCM accumulation on the oocyte nucleus and decreased centrosome microtubule nucleation **a)** Germaria-S5 and S5 control and *vih^ΔN-9^* egg chambers stained for DNA (DAPI) in blue, Plp in green, F-actin (Phalloidin) in red, and centrioles (Asl) in white (asterisks indicate oocytes). **b)** Quantification of Plp staining on the oocyte nucleus S1-3 and germinal vesicle S5 control and *vih^ΔN-9^* egg chambers, significance determined using a Wikcoxon signed-rank test (S1-3: 1×10^−3^<p<5×10^−3^; S5: 1×10^−2^<p<5×10^−1^). **c)** Control and *vih^ΔN-9^* S4/5 egg chambers with the centrosomes marked with the centriole marker Asterless (Asl)-YFP and growing microtubules were labeled with the plus-end marker EB1-GFP (arrows indicate centrioles). All magnification bars equal to 10 μm.

### vih^ΔN-9^ centrosomes have decreased microtubule-nucleating ability

To further test the model that the centrosomes are required for MTOC transfer, we determined if *vih^ΔN-9^* egg chamber centrosomes have decreased microtubule nucleating ability compared to controls. We assessed the microtubule nucleating capacity of both S4-5 control and *vih^ΔN-9^* nurse cell-associated centrosomes live. To avoid pleiotropic effects, we selected *vih^ΔN-9^* egg chambers that did not display the membrane phenotype and still had an oocyte-centric MTOC, as shown by the gradient of microtubule density peaking at the oocyte. In our experimental conditions (see Methods), we confirmed the reduced nucleating ability of *vih^ΔN-9^* centrosomes, 11%, compared to controls, 24% (Fig. 4c and Extended Data Movie 3-4). Additionally, in S4-6 control egg chambers there is one oocyte-centric MTOC (Extended Data Fig. 4a). In contrast, weak bundles of microtubules appeared to grow from the multiple ectopic centrosome/F-actin clusters in all S4-6 *vih^ΔN-9^* egg chambers exhibiting the membrane phenotype (Extended Data Fig. 4a). To further assess the nucleating ability of the *vih^ΔN-9^* centrosomes we looked at Dynein localization. Dynein (Dhc64C), is necessary for the transport of the centrosomes to the oocyte (Bolívar et al., 2001), and it is reported to localize to the minus-end of microtubules at the oocyte MTOC during S4-6 of oogenesis, which is on the periphery of the germinal vesicle (Li et al., 1994). Similar to the decreased Plp staining observed in S1-3 *vih^ΔN-9^* egg chambers from ovarioles with the membrane phenotype in later stage egg chambers, there was also a significant reduction in Dynein localization compared to controls (Extended Data Fig. 4b,c). Furthermore, in support of the microtubule staining characteristic of S4-6 *vih^ΔN-9^* egg chambers harboring the membrane phenotype, Dynein accumulation associated with the centrosomes/F-actin clumps in these egg chambers (Extended Data Fig. 4b).

### Vih is required for the maintenance of oocyte fate

In *vih^ΔN-9^* egg chambers with the membrane phenotype we did not observe a morphologically distinguishable “oocyte” (typically identifiable after stage 3 of oogenesis), which was expected since egg chambers with disrupted microtubule organization do not localize the fate determinants to a single cell and thus oocyte fate is lost or not obtained (Huynh and St Johnston, 2004; Theurkauf et al., 1993). We therefore checked if control of APC/C-Vih activity is required for oocyte fate determination and maintenance. First, we determined if the oocyte was initially selected in *vih^ΔN-9^* egg chambers by staining for Orb, an oocyte fate marker that is localized in S1 (Extended Data Fig. 5a) (Mahowald and Strassheim, 1970). In *vih^ΔN-9^* egg chambers Orb always accumulated more strongly in a single cell in S1 (Extended Data Fig. 5a) and remained localized to a single cell in *vih^ΔN-9^* egg chambers with intact germline cells. However, in 95% of *vih^ΔN-9^* egg chambers displaying the membrane phenotype, Orb failed to remain accumulated in the oocyte after S3, indicating that oocyte fate was subsequently lost, a likely consequence of the membrane rupture (Extended Data Fig. 5a). Second, we checked whether these oocytes were no longer arrested in meiosis by analyzing the expression of the meiotic cell cycle markers Dacapo (Dap) and Cyclin E (CycE) (Huynh and St Johnston, 2004). In control egg chambers, both Dap and CycE localized to the germinal vesicle in the oocyte early during oogenesis and remained there until late oogenesis (Extended Data Fig. 5b). However, in 86% of *vih^ΔN-9^* egg chambers, there was no obvious Dap and CycE staining within the germinal vesicle after S3 (Extended Data Fig. 5b). We conclude that *vih^ΔN-9^* oocytes often overcome the meiotic arrest typical of control oocytes and slip into the endocycle. Together, these results indicate that the oocyte was initially specified, and that the initial fate determination does not require the localisation of the centrosomes, consistent with previously findings (Bolívar et al., 2001; Röper and Brown, 2004)

An immediate consequence of the loss of oocyte fate in *vih^ΔN-9^* egg chambers should be a drop in the overall production of functional eggs and in *vih^ΔN-9^* flies we observed a 55% decrease in cumulative fecundity (total number of eggs laid over a 21-day period) compared to control (Extended Data Fig. 5c). This phenotype worsened with age, similar the increased penetrance of the membrane and centrosome phenotypes in older females (Extended Data Fig. 5d). In combination with the decrease in fertility, we observed a proportion of laid eggs with dorsal appendage defects (9%, n=836), indicative of dorsal-ventral abnormalities, which was not observed in controls (0%, n=283) and is likely the result of oocyte MTOC disruption.

## Concluding Remarks

Syncytial cyst centrosome migration occurs during oogenesis in many insects and even mammals (Lei and Spradling, 2016). In this study, we provide strong evidence that in the *Drosophila* female germline, proper regulation of the stability and microtubule nucleation activity of centrosomes is essential for their effective transportation to the oocyte. Additionally, our data support the hypothesis that the functional significance of centrosome migration is to form the oocyte-centric MTOC through PCM transfer to oocyte nucleus. This may in turn promote the microtubule dependent transport of other materials, such as the Golgi bodies, the organization of which is necessary for proper membrane dynamics (Coutelis and Ephrussi, 2007; Cox, 2003). The overall consequence of centrosome transport failure likely results in the destabilization of the delicate membrane structure in the syncytial cyst, a phenotype also seen when there is ectopic microtubule organization in the egg chamber caused by reduced *γ*-tubulin (a key PCM component) (Tavosanis and Gonzalez, 2003) or ectopic spindle formation caused by APC/C inactivation (Reed and Orr-Weaver, 1997).

Our findings may appear contrary to previously findings that a substantial reduction in centrosome numbers had no phenotype in oocyte development (Stevens et al., 2007). However, our results are a consequence of the centrosomes being misplaced, not eliminated, thereby demonstrating the necessity of their transport within the syncytial cyst. Furthermore, the method used by the previous study for centrosome elimination left some remaining centrosomes which may have been sufficient to complete their normal function. Additionally, our study also highlights the evolutionary conserved, APC/C-Polo module, as a key upstream component of natural centriole elimination, providing a further insight into the molecular mechanism underlying this phenomenon (Lei and Spradling, 2016).

We propose a model where after mitotic exit in the germline cyst, the APC/C must be “turned off” to prevent it from precociously targeting Polo for destruction during a time when Polo is needed to maintain the stable PCM in order to enable centrosome transport to the oocyte. The APC/C then turns back on during a cell cycle transition at ∼S4 (Reed and Orr-Weaver, 1997), at which point the APC/C starts to degrade Polo to enable natural centriole elimination (Pimenta-Marques et al., 2016). Centrosome misregulation is a common occurrence in many diseases, including cancer, and our results extend beyond the basic biology of animal fertility to give a potential indication as to why over-activity of Ube2c is common in many cancerous tissues and indicative of a poor prognosis (Broutier et al., 2017).

## Acknowledgements

We are thankful to Margit Pal for the creation of transgenic lines. We are also thankful to E. Clark, J. Casal, T. Martins, Juan Jose Perez-Moreno, A.T.C Carpenter, and C. Östergren for critical reading of the manuscript. We would also like to thank all members of the Y. Kimata, D.M. Glover, P.A. Lawrence, and I.M. Palacios labs for discussions. We thank D. St Johnston, H. Yamano, N. Rusan, F. Port, J. Raff, and P. Conduit, for sharing tools. We would like to thank the Genetics Fly Facility, University of Cambridge, for embryo injections. Both ALB and FM were funded by Cancer Research UK (CRUK-A12874). Additionally, ALB was funded by The Wellcome Trust Institutional Strategic Support Fund (ISSF) and Cancer Research UK (CRUK-RG78567). The Kimata and Glover Labs are both funded by CRUK.

## Methods

### Drosophila Stocks

All stocks were maintained at 25°C. All CRISPR/Cas9 engineered stocks were created as detailed below. The nos-cas9 and act-cas9 stocks were a kind gift from Fillip Port (Port et al., 2014). The transgenic gRNA flies were created using either *y^1^ sc^1^ v^1^ P{nos-phiC31\int.NLS}X; P{CaryP}attP2* (BDSC 25710) or *y^1^ v^1^ P{nos-phiC31\int.NLS}X; P{CaryP}attP40* (BDSC 25709). The controls used were either *w^1118^* or w^+O^, except for the internal control which was w*; *P{w+^mC^=Ubi-GFP.D}61EF P{w+^mW.hs^=FRT(w^hs^)}2A* (BDSC 1626). The FRT chromosome that the *vih^ΔN-4^* allele was created on was *w* P{w+^mW.hs^=FRT(w^hs^)}2A* (BDSC 1997). The *vih* deficiencies used for the complementation test were *w^1118^; Dƒ(3L)BSC380/TM6C, Sb^1^ cu^1^ (BDSC-24404)* and *w^1118^; Dƒ(3L)ED4483, P{3’.RS5+3.3’}ED4483/TM6C, cu^1^* Sb^1^ (BDSC-8070). The ubiquitously expressed transgenic fly lines we generated for this study: *P{Ubi-vih-WT}, P{Ubi-vih^ΔN-9^}* and *P{Ubi-vih^ΔCS^}.* The *mr* alleles were obtained from the Bloomington Drosophila Stock Center (BDSC), *e(mr)^1^;mr^2^/SM6a* (BDSC-4535) and *px^1^ bw^1^ mr^1^ sp^1^/In(2LR)bw^V1^, ds^33k^ bw^V1^* (BDSC-380) (Reed and Orr-Weaver, 1997). The GFP-Polo stock is expressed using the endogenous promotor *P{GFP-polo.wt}* was a kind gift from Adelaide Carpenter (Moutinho-Santos et al., 1999). The *Ubi-cycB-GFP* was a kind gift Jordan Raff. The *polo^1^* and *polo^11^* alleles were a kind gift from Adelaide Carpenter. The *Sas4^s2214^* allele was a kind gift from Paul Conduit (Basto et al., 2006). The *asl:YFP* allele was a kind gift from C. González (Varmark et al., 2007). The *Eb1:GFP* allele originated from the Uemura laboratory (Shimada et al., 2006).

### gRNA design, expression plasmids, and transgenic fly-lines

The gRNA sequenced used was CGAGCAAAGTGGTGGAGCAG. The target site was selected in order to direct Cas9 cleavage to just after the 5’ start methionine codon of the transcript. The CRISPR Optimal Target Finder (http://tools.flycrispr.molbio.wisc.edu/targetFinder/) was used to design the 20mer target sequence (Gratz et al., 2014). Off-target assessment was done with this program and the target site was chosen because it did not have any homologous sites anywhere else in the genome. The gRNA plasmid pCFD3 (a gift from Fillip Port) was used to create the gRNA transgenic flies (Port et al., 2014). The oligos were cloned directly into the pCFD3 vector for production of the transgenic gRNA line. All fly embryos were injected by the Cambridge Genetic Fly Facility, using standard procedures. The gRNA expression plasmid was inserted into the genome using the Phi31C system at the attP40 site.

### CRISPR/Cas9 Fly genetics

For each experiment, transgenic *cas9* virgin females were crossed to U6-gRNA–expressing males carrying the gRNA plasmid insertion, similar to previous studies (Port et al., 2014). Individual females resulting from this cross were subsequently crossed to a balancer stock to remove the Cas9 expression and the resulting single females were again crossed to the balancer stock to generate a stable fly line. Therefore, the other chromosomes that were present during the mutagenesis were eliminated during stock creation. When the alleles were created in an FRT background, the gRNA line was first balanced and crossed to flies containing the FRT chromosome; males of this line were then crossed to transgenic *cas9* virgin females and carried through the normal procedure.

### Target Loci Characterization

In order to identify CRISPR/Cas9-induced mutations, genomic DNA was isolated from flies by crushing them in 200 μL of BufferA (100 mM Tris pH 9.0, 100mM EDTA, 1% SDS), and incubating the lysate at 65-70°C for 30 min. The lysate was chilled on ice for 5 min, before adding 30 μl of 7.5M NH_4_-acetate. The lysate was then mixed gently, chilled on ice for 20 min and centrifuged for 10 min at 13,000 rpm. The supernatant was transferred to a new tube and spun again for 10 min. The supernatant was again transferred to a new tube, mixed gently with 150 μl of isopropanol, and spun again. The supernatant was removed and the DNA pellet was washed with 70% ethanol. The DNA was then dissolved in 100 μl of dH_2_O. 1 μl was used per PCR reaction. PCR was performed with the primers: AAGCCAAACAGCGGATAGCA and TGGTTGGCAGTAGGTCTTCG. PCR products were gel verified and sent for sequencing. The sequencing of genomic DNA from heterozygous flies yielded an overlay of the DNA sequence from both chromosomes. The mutant sequence was then manually called from the chromatogram.

### CRISPR allele characterization and additional phenotypic data

We specifically targeted the N-terminal region of *vih* with a transgenically expressed gRNA (Fig. 1A). The 27-amino acid N-terminal region is unique to the Ube2c family (Fig. 1B) (Fumio et al., 1997), and has been shown to be necessary for APC/C regulatory function in human cell lines (Summers et al., 2008; Williamson et al., 2009). The rest of the protein consists of the core domain, which contains two destruction boxes plus the UBC domain, which contains a catalytic cysteine residue (Mathe et al., 2004). The core domain is thought to be responsible for the specificity of Vih to the APC/C (Fig. 1C) (Summers et al., 2008). We created two different types of mutations (Fig. 1C): two null mutants (*vih^Null^*), which are frame-shift mutations, and two *vih^ΔN^* alleles with in-frame deletions within the 27-amino acid N-terminal region.

The *vih^Null^* alleles have either 1- or 4-nucleotide deletion, resulting in a frame-shift at the 8th amino acid of the gene, which disrupts all domains of the protein. The *vih^Null^* alleles were created using different Cas9-expressing lines in different genetic backgrounds. None of these alleles are homozygous viable after larval stage 3, and they do not complement two *vih* deficiencies nor the *vih^KG02013^* allele (Nagy et al., 2012). We were able to rescue the viability of the two null alleles with ubiquitously expressed Vih^WT^ and Vih^ΔN-9^ but we were not able to rescue viability with Vih^ΔCS^, a catalytically dead version of Vih, and only the transgenically expressed Vih^WT^ allele was able to rescue the fertility (data not shown).

The *vih^ΔN-9^* mutation is a deletion of 9 amino acids in the 27-amino acid N-terminal region. The *vih^ΔN-9^* allele is homozygous viable and complements two *vih* deficiencies. The other N-terminal allele, *vih^ΔN^* allele (*vih^ΔN-4^*) is a deletion of only 4 amino acids. This allele was created using a different Cas9-expressing line directly on an FRT chromosome, and was therefore created in a completely different genetic background. Both *vih^ΔN^* alleles delete part of the Ube2c family specific 27-amino acid N-terminal region (Fumio et al., 1997). Transheterozygous *vih^ΔN-9^/ vih^ΔN-4^* has the same phenotype as the *vih^ΔN-4^* allele alone and neither alleles have a phenotype when crossed to a *vih* deficiency (Data not shown). The phenotype that was present in both alleles increased with age and an age of 14 days was chosen for analysis, unless otherwise stated. The 3rd chromosome with the *vih^ΔN-9^* allele on it was isogenised with *w^+O^* after the stock was originally created and rebalanced.

### Western Blots

For the western blots of *Drosophila* ovaries, 30 ovaries were dissected in 60 μl of PBS containing Protease and Phosphatase inhibitor (Roche). The samples were lysed using a homogenizing pestle (Sigma-Aldrich), and the lysates were clarified by centrifugation. 2x Laemmli buffer was added to the cleared lysates, and the samples were boiled 2 min. The proteins were then resolved by SDS-PAGE. The electrophoretic run was performed using a Mini-PROTEAN Tetra Cell System (BioRad), in a Running Buffer solution (25 mM Tris, 192 mM glycine and 0.1% SDS, pH approx. 8.6, Sigma), at 200 Volts (V). The gel was assembled in a ‘transfer sandwich’ (cushion pad-filter paper-gel-membrane-filter paper-cushion pad) and blotted on a nitrocellulose membrane (GE Healthcare) for 2 hr at 60 V in Transfer Buffer solution (25 mM Tris, 190 mM Glycine, 20% Methanol). Protein transfer was verified by Ponceau S staining, and the membrane was incubated for 45 min at room temperature in a blocking solution containing 5% milk (Marvel) and 0.1% Tween (Sigma) in PBS. The membrane was then incubated in a primary antibody solution prepared in blocking solution for 1 hour at room temperature. The membrane was washed three times for 10 min at room temperature with a solution of 0.1% Tween in PBS (PBST) and then incubated in a secondary antibody solution prepared in blocking solution for 1 hour at room temperature. The membrane was washed three times with PBST for 10 min at room temperature, incubated with a peroxidase ECL substrate (Pierce), and the proteins were detected by exposing an X-ray film (Fuji). **Primary antibodies:** Mouse anti-α-tubulin (1:1000) Clone DM1A from Sigma, Mouse α-Polo (1:500) and Rabbit α-Vihar (1:500), Glover Lab.

### Egg-lay and hatching rate

In large cages, 50 virgin females were mated with equal numbers males and maintained on apple juice agar plates with fresh yeast; the cages were kept at 25°C. The number of eggs laid during 2 × 2 hour collections were counted every 1-3 days for a 21 or 28-day period (N=2). The number of flies in the cadges were kept equal throughout the course of the experiment, the number of female flies did not drop below 40. After the eggs were counted the plates were kept at 25°C for 5 extra days and examined for the number of larvae hatched. The hatch-rates were calculated by dividing the number of larvae that eclosed by the total number of eggs laid.

### Dissection and immunofluorescence

Fixed sample preparation: After fattening 14-day aged and mated females that were maintained at 25°C for 16-24 hours with extra yeast, ovaries were dissected in 0.2% PBT (PBS + 0.2% Tween), fixed for 20 min in 4% paraformaldehyde/PBT, washed with PBT, blocked with PBT + 10% BSA for 1 hour, and incubated with the primary antibody in PBS + 2% Tween + 1% BSA for 16-24 hours at 4°C. After washing the ovaries with PBT three times for a total of 25 min, they were incubated with the secondary antibody for 2 hours at RT, or 16 hours at 4°C. Phalloidin staining for visualizing F-actin was done for 20 min, either after fixing or after secondary antibody staining. Finally, the ovaries were washed twice with PBT for a total of 40 min and mounted in Vectashield (Vector) with DAPI for visualizing DNA. The microtubule sample preparation and staining was performed as previously published (Legent et al., 2015). Unless specified, all steps were performed at room temperature. **Internal Controls:** GFP containing flies were used and taken through the whole procedure (from fattening to imaging) mixed with the mutant samples to ensure that the membrane phenotype was not due to physical damage. **Primary antibodies:** Mouse α-Orb 4H8 and 6H4 (1:100 each) from Developmental Studies Hybridoma Bank (DSHB), was a kind gift from Daniel St Johnston. Guinea Pig α-Asterless (1:40,000) was a kind gift from Nasser Rusan (Lerit and Rusan, 2013). Mouse anti-α-tubulin (1:100) Clone B-5-1-2 from Sigma. Mouse α-Dap NP1-s (1:10) from DSHB. Rabbit α-Cyclin E (1:100) from Santa Cruz Biotechnology. Mouse α-Hts 1B1-s (1:30) from DSHB. Mouse α-Polo (1:50), Rabbit α- Vihar (1:500), Mouse α-Dynein IC74 (1:1000), and Chicken D-Plp (1:1000), Glover Lab. **Secondary antibodies:** (all 1:500) Goat α- Chicken 488 from Life Technologies, Goat α- Mouse 488 and 647 from Life Technologies, Goat α-Guinea Pig 488 from Invitrogen and 647 from Life Technologies, Goat α-Rabbit 488 from Invitrogen and 647 from Life Technologies. **Small molecule stains:** Alexa Fluor™ 568 Phalloidin (1:200) from Thermo Fisher Scientific, CellMask™ Deep Red Plasma Membrane Stain (1:1000) from Thermo Fisher Scientific.

### Intensity measurements of fixed samples

For the Plp and Dynein experiments, the control and *vih^ΔN-9^* egg chamber samples were prepared in parallel and treated the same. The control was analyzed first to establish the confocal settings to be used, which remained unaltered for all images acquired, for the channel being directly compared (which was the 488 channel for all experiments). The intensity measurement was taken using FIJI measurement tool. The measurement section chosen for the Plp and Dynein was a box drawn to the edges of the nucleus at early time points and germinal vesicle for S4 and later. Special care was taken to not include the centrosome associated Plp or Dynein fluorescence in the intensity measurement analysis, since they are closely associated with the oocyte nucleus and germinal vesicles in the control but not *vih^ΔN-9^* egg chambers. Stacks of egg chambers were taken every 1μm and the brightest single image was chosen for the intensity measurement analysis. The relative intensity was calculated by dividing by the average of the control.

### Live sample preparation

After fattening, 14 day aged mated females that were maintained at 25°C for 16-24 hours were dissected and their ovaries were short-term live imaged (1 hour at room temperature) by dissecting directly in voltelef oil or in PBS with CellMask™ (incubated for 15 min prior to being transferred to voltelef oil for imaging).

### Microtubule nucleation live-imaging sample preparation

Ovaries from flies aged for 14 days were dissected for in vivo preparation as previously described (Diaz de la Loza et al., 2017; Valencia-Exposito et al., 2016). Images were acquired with a ZEISS LSM 880 Airyscan confocal microscope and analyzed using ImageJ. In vivo images were obtained at ∼25 °C with a 40×/1.2 numerical aperture (NA) water immersion objective. Single-focal planes were taken every 2 seconds and processed with proprietary ZEISS’ ZEN 2.1 software for Airyscan resolution optimization.

### Ubiquitination assay

For the ubiquitination assay, *Drosophila* APC/C was immuneprecipitated from 2 grams of pMTB-Apc4-TAP *Drosophila* embryos or from 0.5–1×109 of either pMT-PrA-Cdc27 or pMT-Apc4-TAP D.mel2 cell lines. The samples were resuspended in 10 ml of ice-cold Buffer A (75 mM HEPES pH7.5, 150 mM KCl, 1.5 mM EGTA, 1.5 mM MgCl2 a, 7.5% glycerol, 0.1% NP40) containing fresh DTT (5 mM) and complete protease inhibitor cocktail (Roche). Embryos were broken using a glass homogenizer and the lysates were cleared by centrifugation at 20,000 rpm for 30 min. The cleared lysates were mixed with preequilibrated Dynabeads (Invitrogen) conjugated with rabbit IgG (MP Biochemicals) and incubated for 2–4 hours with gentle rotation. Non-specific bound proteins were removed by six successive washes in Buffer A which contained low salt followed by a final wash with Buffer B (50 mM Tris-HCl pH 8.0, 0.5 mM EDTA). The E2 enzymes His-Vih^WT^, His-Vih^ΔN-9^, His-Vih^ΔCS^ and His-Ube2s were expressed in *E. coli* (strain Bl21 cod+) by incubating the cultures overnight at 18°C with 0,1 mM of IPTG. The fusion proteins were then purified using Ni-NTA Agarose beads (QIAGEN). Substrates were labeled with [35S] methionine (PerkinElmer) in a coupled in vitro transcription-translation (IVT) system (Promega). His-Fzy was purified from insect cells using the baculovirus-based expression system. Briefly, a plasmid containing His-Fzy was transformed into MAX Efficiency DH10Bac competent cells (Thermo Fisher Scientific) and the bacmids carrying His-Fzy genes were purified according to manufacturer’s instructions. The bacmids were handed to the Baculovirus Facility at the Department of Biochemistry of the University of Cambridge for insect cells transfection, virus production and titration. His-Fzy viruses were transduced into Sf9 cells at a MOI of 10. His-Fzy was purified from cell pellets from 800 mL of liquid culture of Sf9 cells using Ni-NTA agarose (QIAGEN), according to the manufacturer’s instructions. Ubiquitination reactions were performed at 27°C in 10 μl of the buffer (20 mM Tris-HCl [pH 7.5], 100 mM KCl, 2.5 mM MgCl2) containing 4 μl of purified APC/C, 3.5 μM of E2 enzymes, 0.75 mg/ml ubiquitin, 1 μM ubiquitin-aldehyde, 200 μM MG132, 200μM DTT, 2 mM ATP, and 1 μl in vitro-translated (IVT) Mes1 substrate (Kimata et al., 2008). Reactions were stopped at the indicated time points with SDS sample buffer and mixtures were resolved by SDS-PAGE.

### Destruction assay

The in vitro destruction assay in *Xenopus* egg extracts was performed as described before (Yamano et al., 2009). Briefly, ^35^S-methionine-labeled substrate proteins were prepared in a coupled in vitro transcription-translation system (Promega) according to manufacturer’s instructions. Cytoplasmic extracts of cytostatic factor (CSF)-arrested *Xenopus* eggs were prepared following standard procedures. The CSF extracts were first released into interphase by addition of 0.4 mM CaCl_2_ and 10 μg ml^-1^ cycloheximide, and then incubated for 2-4 hours at 23 °C. Substrates were added to the extracts and the reactions were started by adding 150 ng of purified *Xenopus* Fzr (gift from Hiro Yamano) to the mixture. Aliquots were collected into 2x Laemmli buffer at 0, 1, 2 and 3 hours, boiled for 2 min and resolved by SDS-PAGE.

### Statistical Test

z-test for proportions: the n-values are given on the figures and the p-values are given in the figure legends. Shapiro–Wilk for normalcy: all non-proportional datasets were tested for normalcy and all came back with skewed bell curve distributions (not normal). Wikcoxon signed-rank test for under 20 samples (given as a range because it is an estimate with small sample sizes): the n-values are given on the figures and the p-value ranges are given in the figure legends. Welch unbiased two-tailed t-test used for non-normal sample sizes of 20 or greater: the n-values are given on the figures and the p-values are given in the figure legends. When this test was used the non-parametric Wikcoxon signed-rank test was also performed and the p-values obtained fell within the range given by the Welch unbiased two-tailed t-test. This was used to give a specific p-value when possible.

